# Detecting turnover among complex communities using null models: A case study with sky-island haemosporidian parasites

**DOI:** 10.1101/2020.04.17.046631

**Authors:** Lisa N. Barrow, Selina M. Bauernfeind, Paxton A. Cruz, Jessie L. Williamson, Daniele L. Wiley, John E. Ford, Matthew J. Baumann, Serina S. Brady, Andrea N. Chavez, Chauncey R. Gadek, Spencer C. Galen, Andrew B. Johnson, Xena M. Mapel, Rosario A. Marroquin-Flores, Taylor E. Martinez, Jenna M. McCullough, Jade E. McLaughlin, Christopher C. Witt

## Abstract

Turnover in species composition between sites, or beta diversity, is a critical component of species diversity that is typically influenced by geography, environment, and biotic interactions. Quantifying turnover is particularly challenging, however, in multi-host, multi-parasite assemblages where undersampling is unavoidable, resulting in inflated estimates of turnover and uncertainty about its spatial scale. We developed and implemented a framework using null models to test for community turnover in avian haemosporidian communities of three sky islands in the southwestern United States. We screened 776 birds for haemosporidian parasites from three genera (*Parahaemoproteus, Plasmodium*, and *Leucocytozoon*) by amplifying and sequencing a mitochondrial DNA barcode. We detected infections in 280 birds (36.1%), sequenced 357 infections, and found a total of 99 parasite haplotypes. When compared to communities simulated from a regional pool, we observed more unique, single-mountain haplotypes and fewer haplotypes shared among three mountain ranges than expected, indicating that haemosporidian communities differ to some degree among adjacent mountain ranges. These results were robust even after pruning datasets to include only identical sets of host species, and they were consistent for two of the three haemosporidian genera. The two more distant mountain ranges were more similar to each other than the one located centrally, suggesting that the differences we detected were due to stochastic colonization-extirpation dynamics. These results demonstrate that avian haemosporidian communities of temperate-zone forests differ on relatively fine spatial scales associated with adjacent sky-islands. Null models are essential tools for detecting turnover in complex, undersampled, and poorly known systems.

## Introduction

The mechanisms that cause community composition to change across space are critical to the origin and maintenance of species diversity, and are therefore worthy of detailed study (Buckley and Jetz 2008; Socolar et al. 2016). Species diversity is traditionally partitioned into local (alpha) and regional (gamma) scales, with beta diversity, defined as species turnover between local communities, representing a link between local and regional processes (Whittaker 1960, 1972). Beta diversity is affected by geographic distance, environmental factors (abiotic), and species interactions (biotic), and considerable efforts have been made to disentangle the roles of these various factors in determining community composition and structure (e.g., Fitzpatrick et al. 2013, Warburton et al. 2016, Clark et al. 2018). A major challenge for quantifying turnover, however, is the inability to fully sample the communities being compared. Sampling artifacts can either inflate or diminish beta diversity estimates because of undersampling, differences in abundance or detectability of species, or the proportion of rare species in a given system (Colwell and Coddington 1994; Chao et al. 2005; Cardoso et al. 2009). The issues of undersampling or biased sampling are especially complex in multi-host, multi-parasite assemblages in which parasite communities are nested within host communities (Poulin 1997; Krasnov et al. 2011), hosts and parasites have reciprocal filtering effects (Poulin 1999; Brooks et al. 2006; Thornhill and Fincher 2013), and undersampling of rare or host-specific microbial parasites is unavoidable (Hughes et al. 2001; Zhou et al. 2013; Poulin 2014).

Several studies have called attention to sampling effects on beta diversity estimation and have taken various approaches to address this issue. One suggested approach was to develop new beta diversity metrics that adjust for unsampled species (Chao et al. 2005, 2006), although these metrics still appear to be sensitive to sampling under certain conditions (Beck et al. 2013). Other studies have attempted to identify beta diversity metrics that are less sensitive to sampling effects, in order to provide recommendations for which metrics are most appropriate to use for a given question and system (Cardoso et al. 2009; Beck et al. 2013). For example, the *ß*_–2_ (Harrison et al. 1992) and *ß*_–3_ (Williams 1996) metrics were found to be particularly robust to undersampling but were insensitive to species richness differences between communities, such that other metrics may be preferred if richness differences are important (Cardoso et al. 2009). Rarefaction techniques, which adjust for sampling effort through random subsampling, are well-established for alpha diversity and have more recently been extended to estimate beta diversity (Dornelas et al. 2014; Stier et al. 2016). These methods can account for uneven sampling among sites and thus produce comparable relative estimates of beta diversity, but at the inevitable potential cost of removing data and statistical power. By reducing sampling, these approaches can lead to the appearance of community differences, leaving uncertainty about whether there is turnover or not.

Null models have also been highlighted as a useful tool for studying beta diversity that could be used to address sampling issues (Anderson et al. 2011). For example, Roden et al. (2018) used null models to demonstrate conditions under which beta diversity can be accurately estimated by focusing only on abundant taxa. Their conclusions rest on the assumption that patterns evident in dominant taxa can be extrapolated to the whole community; however, this may not be a suitable assumption for diverse pathogen or parasite communities in which rarer taxa can have ecological significance due to varying degrees of host-specialization and phenomena such as periodic host-switching (Brooks and Hoberg 2007; Woolhouse and Gaunt 2007; Ricklefs et al. 2014). Ideally, community differences in complex host-parasite assemblages should be tested with a method that accounts for undersampling of parasites, uneven and incomplete sampling of hosts, and substantial variation in abundance and host specialization among parasite species.

Here, we implement a null modeling framework to test for and investigate community turnover in the multi-host, multi-parasite assemblage of haemosporidian parasites and their avian hosts in the sky islands of the southwestern United States. These sky islands are defined by high-elevation forested habitats that are geographically isolated by low-elevation, arid habitats. Similar to archipelago systems (Ricklefs and Bermingham 2008), sky islands provide opportunities to study the factors that influence community structure, biogeographic patterns, and evolutionary diversification (Knowles 2001; McCormack et al. 2008; Gupta et al. 2019; Williamson et al. 2019). Avian haemosporidians include the intracellular, protozoan parasites that cause avian malaria, a global disease system that has been associated with epidemics, population declines, and extinction of naïve hosts (Warner 1968; van Riper et al. 1986; Atkinson and LaPointe 2009). These remarkably complex assemblages are becoming model systems for studying ecology and evolution of host-parasite interactions. At least three genera of haemosporidians (*Plasmodium, Parahaemoproteus*, and *Leucocytozoon*) commonly infect diverse bird communities with varying degrees of host specificity (Hellgren et al. 2009; Loiseau et al. 2012), abundance (Fallon et al. 2005; Ricklefs et al. 2011), and potential impacts on survival and fitness (Marzal et al. 2008; Knowles et al. 2010; Asghar et al. 2015). The geographic scale and potential drivers of haemosporidian community turnover have been investigated in several continental and island systems (e.g., Scordato and Kardish 2014, Ellis et al. 2015, Clark and Clegg 2017), but few consistent patterns have emerged across regions, host species, or parasite genera, underscoring the complexity of these systems.

On a global scale, haemosporidian communities do not appear to follow common patterns of biodiversity such as latitudinal gradients (Clark 2018; Fecchio et al. 2020). On a regional scale, host community turnover has been one of the most consistent predictors of haemosporidian community turnover across systems including Melanesia (Olsson-Pons et al. 2015; Clark et al. 2018), South America (Fecchio et al. 2017), eastern North America (Ellis et al. 2015), and the Southwestern U.S. (Williamson et al. 2019). In contrast, geographic distance has been implicated in parasite community dissimilarity in some regions (West Indies: Fallon et al. 2005; South America: Fecchio et al. 2017), but not others (Asia: Scordato and Kardish 2014; Southwestern U.S.: Williamson et al. 2019). Distance-decay patterns also vary among haemosporidian genera, with *Plasmodium* exhibiting more geographic structure than *Parahaemoproteus* in two island systems (Fallon et al. 2003; Ishtiaq et al. 2010), despite the common finding that *Plasmodium* lineages tend to be more generalized and can be globally distributed (Hellgren et al. 2015; Walther et al. 2016). Similarly, environmental characteristics such as elevation and climate influence the distribution of haemosporidian genera in different ways, a finding that may be related to variation in the distribution of genus-specific vectors (Valkiu□nas 2005; van Rooyen et al. 2013; Galen and Witt 2014; Harrigan et al. 2014). In sum, disentangling the drivers of haemosporidian distributions is an exciting and active area of research, but one that is complicated by the effects of chronic undersampling.

Although sampling issues are unavoidable in avian haemosporidian studies, the influence of undersampling on estimates of community turnover have not been explicitly addressed. Instead, authors have taken pragmatic approaches such as including only well-sampled species or lineages in their analyses (Ellis et al. 2015; Soares et al. 2017), combining sites into regional communities for analysis (Clark 2018), choosing beta diversity metrics that are thought to be less sensitive to sampling issues (Svensson-Coelho and Ricklefs 2011), or using rarefaction methods to evenly sample communities before estimating beta diversity metrics (Fecchio et al. 2019; Williamson et al. 2019). These and other previous studies of turnover were focused on obtaining comparable relative measures of beta diversity to test for its causes, but they may have neglected to address a more fundamental question: Are the communities different at all? This question is important to address with rigor because undersampling of parasite species or uneven sampling of host species may cause absolute estimates of beta diversity to exceed their true values.

The primary goal of our study was to test for turnover among haemosporidian parasite communities of three sky islands in northern New Mexico that are separated by >25 km of unforested habitat and that have near-identical host communities. We compared patterns among the full haemosporidian community of each sky island and, separately, the subsets of the communities comprised of each of the three parasite genera. We estimated the probability that species composition between communities differed after accounting for variation in abundance among parasite species and variation in sampling effort among sky islands. We then conducted four follow-up analyses to better understand our results. First, we assessed sensitivity of results to the host species sampled by repeating tests with only host species that were sampled from all three mountain ranges. Second, we assessed differences in turnover between specialist and generalist parasite lineages. Third, we assessed turnover among sky-island parasite communities of resident host species and migrant host species, respectively. Finally, we assessed turnover among sky island parasite communities within focal host species. While our surveys revealed many parasite haplotypes that occurred on only one or two sky islands, our null model approach was designed to test whether those lineages merely appeared to be restricted due to undersampling alone.

## Materials and methods

### Sampling and parasite identification

We sampled the breeding bird communities from three mountain ranges in northern New Mexico (Zuni Mountains, Mt. Taylor, and Jemez Mountains), as described in Marroquin-Flores *et al*. (2017). We conducted fieldwork between June and July 2016 (n = 186 samples) and 2017 (n = 590 samples) in the transition zone between piñon-juniper woodland and ponderosa pine forests (elevation ∼2100–2500 meters; Fig. 1). Birds were collected by mist-net or shotgun, preserved on dry ice, and transported to the Museum of Southwestern Biology (MSB) at the University of New Mexico for specimen preparation. Thin blood smears were prepared at the time of collection. In 2016, tissues (heart, pectoral muscle, liver) were preserved during specimen preparation at the MSB, and in 2017, tissues and whole blood were flash frozen in liquid nitrogen while in the field. All tissues are cryopreserved in liquid nitrogen at the MSB Division of Genomic Resources. Complete details for each specimen are available in Table A1 (Online Resource 1) and its embedded links to the Arctos database. All samples were collected in accordance with animal care protocols and appropriate state and federal scientific collecting permits.

**Figure 1.**
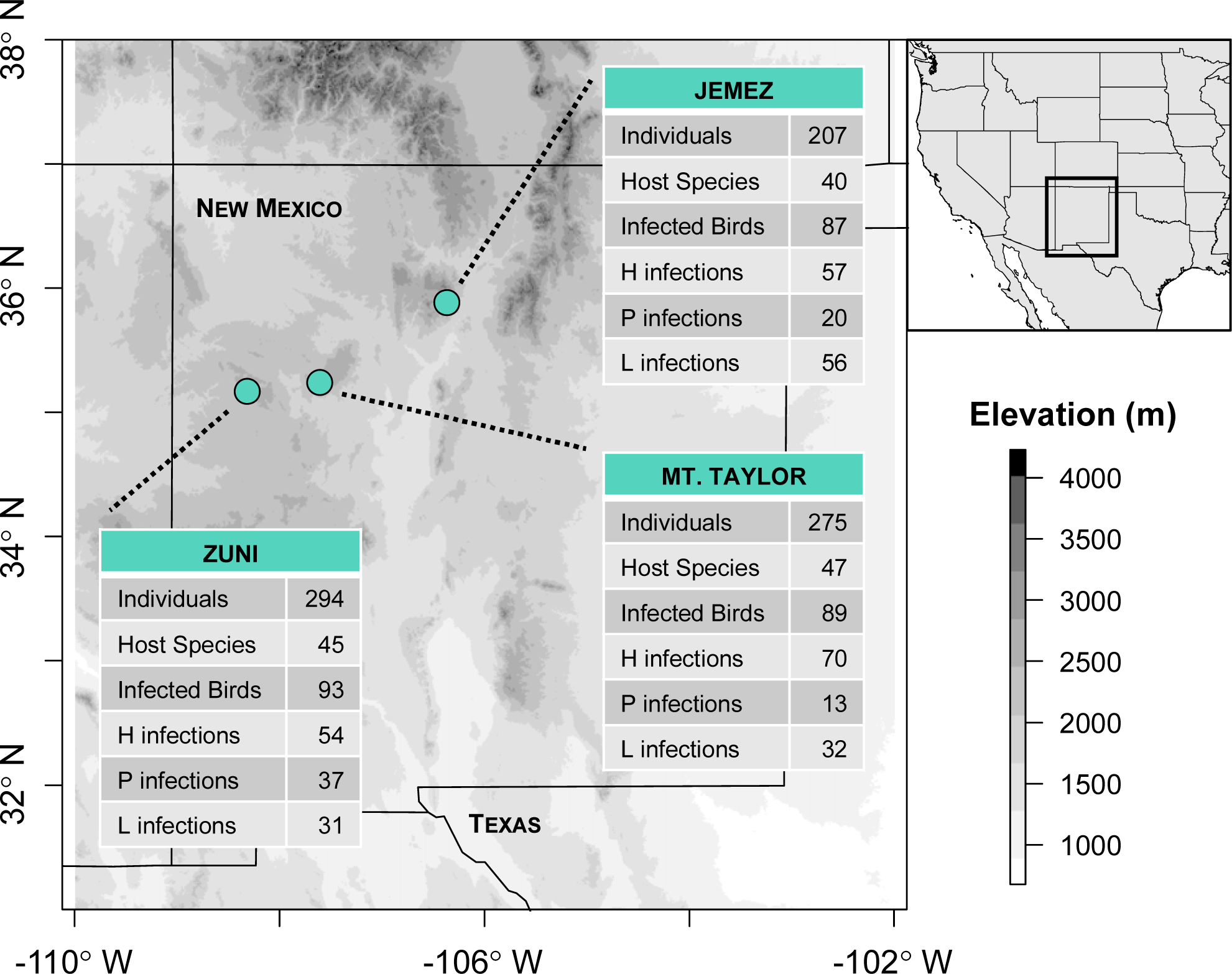
Map of study areas and mountain ranges sampled. The number of individual birds sampled, host species sampled, birds infected, and infections detected using molecular methods are shown for each mountain. H = *Parahaemoproteus*, P = *Plasmodium*, L = *Leucocytozoon*. Elevation is based on the SRTM Digital Elevation Database (Jarvis et al. 2008).

We extracted genomic DNA from 776 bird specimens, primarily from pectoral muscle, using QIAGEN DNeasy Blood and Tissue kits following the manufacturer’s protocols. To identify haemosporidians from three genera (*Parahaemoproteus, Plasmodium, Leucocytozoon*), we used three nested PCR protocols to amplify 478 base pairs of the parasite mitochondrial cytochrome *b* (cytb) gene (Hellgren et al. 2004; Waldenström et al. 2004). Reaction conditions and thermal profiles are described in Marroquin-Flores *et al*. (2017). We visualized reaction products on agarose gels to identify positive samples. We purified amplified products using ExoSap-IT and sequenced them in both directions using BigDye v3.1 terminator cycle sequencing and an ABI 3130 sequencer. Sequences were edited and assembled using Geneious v8.0 (https://www.geneious.com). We compared each sequenced infection to the MalAvi database (Bensch et al. 2009) using BLAST to determine the parasite genus. Then we either assigned known haplotype names for 100% matches or characterized parasite haplotypes as novel. We assigned names to novel haplotypes following MalAvi naming conventions, using the first three letters of both the genus and species of the first host species from which the haplotype was sequenced, followed by a number to denote multiple haplotypes from that host species (e.g., SPIPAS02 for the second haplotype sequenced in *Spizella passerina*). Parasite sequences are archived on MalAvi and GenBank (Accession numbers MF077648–MF077690; MK216024– MK216290).

Microscopic examination of thin blood smears was conducted for 64% of samples, including all samples with positive PCR infections and for a portion of uninfected samples (n = 494 total screened). Blood smears were air dried in the field, fixed in absolute methanol or ethanol, and stained for 50 min with Giemsa solution (pH 7.0). We examined blood smears using light microscopy, first scanning 10,000 erythrocytes at 200–400× magnification to locate and count *Leucocytozoon* infections. We then scanned 10,000 erythrocytes at 1000× magnification with an oil immersion lens to locate and count *Parahaemoproteus* and *Plasmodium* infections and take digital photographs of representative haemosporidians to archive in the Arctos database. We estimated infection intensity, or parasitemia, as the proportion of infected red blood cells out of 10,000 screened.

### Phylogenetic analysis and host specificity

We estimated phylogenetic relationships among unique haemosporidian haplotypes using BEAST 2.4.8 (Bouckaert et al. 2014). To select an appropriate model of nucleotide substitution, we used PartitionFinder2 (Lanfear et al. 2017) with a search of all BEAST models and AICc model selection. We conducted BEAST analyses using the GTR+I+G model with estimated base frequencies, a relaxed lognormal clock, and two independent MCMC runs of 100 million generations each, sampling every 10,000 generations. We checked for convergence using Tracer v. 1.6 (Rambaut et al. 2018), confirming that ESS values for likelihoods and the majority of parameters were >1000. We then combined runs with LogCombiner using 10% burn-in, at which point stationarity was reached, and resampled states every 20,000 generations. We generated the maximum clade credibility (MCC) tree in TreeAnnotator from 9,002 posterior trees. Trees were rooted with the *Leucocytozoon* clade based on recent phylogenetic hypotheses for avian haemosporidians (Borner et al. 2016; Galen et al. 2018).

We obtained a phylogenetic hypothesis for the avian host species in our study from BirdTree.org, choosing the ‘Hackett all species’ set of trees (Hackett et al. 2008; Jetz et al. 2012). To calculate indices of host specificity, we used the ‘picante’ package in R (Kembel et al. 2010; R Core Team 2016). We report both a simple host specificity index, i.e., the total number of host species utilized, and a more complex index that incorporates host phylogenetic relationships and parasite abundance among hosts. The latter index is the weighted mean pairwise distance (MPD) developed for community phylogenetics (Webb et al. 2002) and has been applied previously to avian haemosporidians (e.g., Fallon et al. 2005, Svensson-Coelho et al. 2013). In brief, MPD describes the evolutionary diversity of hosts utilized by each parasite. To account for differences in sampling across parasites, we also calculated the standardized effect size of MPD (SES_MPD_) using the null modeling procedure in ‘picante’ with the independent swap algorithm (Gotelli 2000), 1000 iterations, and 999 randomly generated host-parasite matrices.

We interpreted parasite lineages with negative values of SES_MPD_ (mpd.obs.z) and low p-values (mpd.obs.p < 0.05) as host specialists, while parasites with the highest values of SES_MPD_ were considered to be host generalists. MPD could not be calculated for parasites sampled from a single host species, but we determined the minimum sample size needed to reject the hypothesis that a parasite is a host generalist, following Svensson-Coelho et al. (2013). Briefly, we determined that the probability of detecting a generalist parasite four or more times from a single host, and only that host, is extremely unlikely (p=0.043), as follows. The best-sampled lineage with the highest skew in frequency in different host species was COLBF21 (10 of 22 infections were detected from the host species *Vireo plumbeus*, or 0.455). The probability that four random samples of this generalist parasite would occur in this same host is 0.455^4^ = 0.043, thus if we detected a parasite four or more times from one host species, we classified it as a host specialist.

### Parasite community turnover

We investigated haemosporidian community turnover among mountain ranges using a null modeling approach. We compared our observed sky-island communities to simulated communities expected under a model of no turnover, in which parasite communities in each mountain range are randomly sampled from the same hypothetical regional pool, with parasite richness (total number of haplotypes) and abundance distributions (frequency of each haplotype) based on empirical values derived from the combined data. Using the same framework, we tested for community differences using: 1) the full host-parasite dataset, 2) each parasite genus (*Parahaemoproteus, Plasmodium, Leucocytozoon*) separately, 3) pruned datasets for the full and genus-level data including the set of identical host species sampled from all three mountain ranges (n = 26 host species), 4) parasites classified as either host specialists or host generalists, 5) parasites of either migratory or resident host species, and 6) separate datasets for individual host species that were well-sampled (n > 20 individuals) and included at least three parasite haplotypes from each mountain range. The modeling procedure, described below, was written as a set of functions in R and is available as electronic supplementary material (Online Resource 3).

#### Observed sky-island communities

The input file consisted of a data frame with all sequenced infections as rows, including the locality (mountain range), parasite genus, and haplotype name for each infection. These data were converted into two matrices of observed haplotypes (rows) per mountain range (columns); the first with abundances, or the number of infections observed (as in Fig. 2, right-most columns), and the second converted to a binary matrix that represents presence (1) or absence (0) of each haplotype in each mountain.

**Figure 2.**
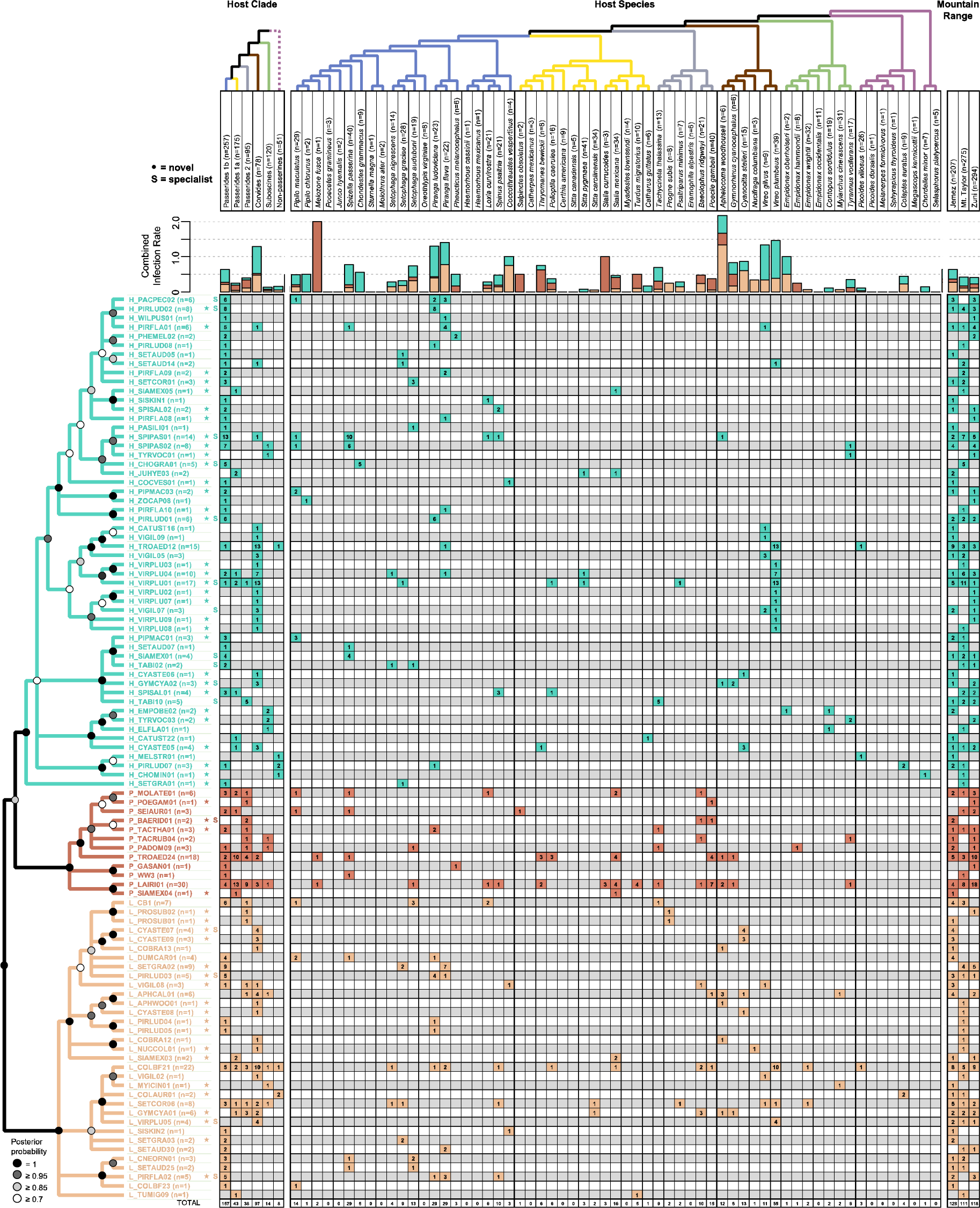
Phylogenetic relationships of haemosporidian haplotypes found in New Mexico birds. Columns represent host clades (left), host species (center), and mountain ranges (right). Dotted line indicates non-monophyly of non-passerines. Host species phylogeny was generated from http://BirdTree.org and branch colors correspond with host clades. Parasite phylogeny was estimated in BEAST and branches with posterior probabilities less than 0.7 were collapsed for visualization. Stars indicate novel haemosporidian haplotypes. “S” indicates haplotypes classified as specialists. Bar plots depict the combined infection rate (number of infections divided by number of birds screened). The number of infections sequenced for each haplotype in each host clade, host species, or mountain range is shown within matrix cells.

#### Hypothetical regional community

We combined the three observed communities and extrapolated to the total expected haplotype diversity using EstimateS v. 9.1.0 (Colwell et al. 2012; Colwell 2013), while assuming that all three mountain ranges are part of a uniform regional community. We created a hypothetical list of haplotypes in the regional community, using the extrapolated total haplotype diversity as a starting point for parasite richness, and with an abundance distribution matching that of the observed communities. For example, if the original community of 99 haplotypes included 33 haplotypes (one-third) with an abundance of one, i.e., each haplotype was sequenced only once, then the hypothetical regional community with extrapolated diversity would also include one-third haplotypes with an abundance of one.

#### Simulated sky-island communities

We generated haplotype lists for the three mountain ranges by stochastically sampling infections from the hypothetical regional community, with replacement, up to the number of observed infections in each mountain range. The probability of sampling each haplotype was weighted by its abundance. We maintained equal richness between simulated and observed sky-island communities by keeping the first 10,000 simulated haplotype lists (i.e. simulated communities) for which the combined number of haplotypes matched that of the observed communities (n=99) once the target number of infections for each of the three mountain ranges had been reached.

#### Test for non-random distribution of haplotype range sizes in the observed data

In order to describe the distribution of range sizes for the simulated communities, we summed the number of haplotypes occurring in a single mountain, shared between two mountains, or occurring in all three mountains, respectively. We compared these sums for the observed data to the distribution of sums from the simulations to test whether, for example, there was an excess of small-ranged (single mountain) haplotypes observed. We calculated p-values as the proportion of simulations in which the number of haplotypes with a given range size (i.e., one, two, or three mountains) was greater than or equal to, or less than or equal to, respectively, the observed number of haplotypes with that range size.

#### Test for non-random dissimilarity among mountain ranges

For each pair of mountains, we calculated the Jaccard dissimilarity index for each simulation and compared the distribution of simulated values to the observed Jaccard values. We used the vegdist() function in the R package “vegan” (Oksanen et al. 2018). Jaccard indices provide a measure of dissimilarity on a scale from 0 (communities identical) to 1 (communities dissimilar). We calculated p-values as the proportion of simulations that were greater than or equal to, or less than or equal to, respectively, the observed Jaccard indices between mountain pairs.

As an additional validation exercise, we compared the abundance distribution of range-restricted haplotypes between simulated and observed communities. In the simulated communities, range-restricted (single mountain range) haplotypes should be dominated by rare haplotypes because all of the haplotypes are assumed to be distributed throughout the region, i.e., they come from the same regional pool. In the observed communities, at least some range-restricted haplotypes should have higher abundances while being truly range restricted. Accordingly, we predict that a cumulative frequency distribution (CFD) of range-restricted haplotypes grouped by abundance (haplotype count) should be right-shifted for the observed community relative to the simulated ones.

## Results

### Parasite abundance and diversity

We detected a total of 280 infected birds out of the 776 that were screened (overall prevalence of 36.1%). We observed parasites broadly across the avian community: 18 of 22 families and 43 of 61 species were infected with haemosporidians. Based on PCR and sequencing results, 155 birds (20.0%) were infected with *Parahaemoproteus*, 62 (8.0%) with *Plasmodium*, and 105 (13.5%) with *Leucocytozoon*. Ninety birds (11.6%) were co-infected with two or more parasite haplotypes. Microscopic examination confirmed positive infections for 168 birds and identified 11 additional infected birds that were not PCR-positive. The highest infection intensity detected was in a sample of *Aphelocoma woodhousii*, which had parasitemia values of 4.8% for *Leucocytozoon* (sequences matched COLBF21/GYMCYA01) and 2.3% of *Parahaemoproteus* (sequence matched SPIPAS01).

Abundances of the three parasite genera differed somewhat among the three mountain ranges (Fig. 1). The highest overall prevalence (proportion of infected individuals from the total screened) was detected in the Jemez Mountains (42.0%), and Mt. Taylor and the Zuni Mountains were similar in prevalence, 32.4% and 31.6%, respectively. *Parahaemoproteus* was the most abundant genus in each of the mountain ranges, followed closely by *Leucocytozoon* in Jemez. *Plasmodium* was more abundant in the Zuni Mountains compared to the other two mountain ranges, where it was the least abundant parasite genus. Host clades and species also varied in prevalence (Fig. 2), with the lowest prevalence in suboscine passerines (10.8%) and the highest in Corvides (74.4%). The species with the highest overall prevalence was *Vireo plumbeus*, with 87.2% of individuals infected.

A total of 357 infections were sequenced and assigned to 99 total haplotypes, including 54 *Parahaemoproteus*, 12 *Plasmodium*, and 33 *Leucocytozoon* haplotypes. When compared to the MalAvi database, 55 (more than half, or a total of 56%) of these haplotypes were newly described from New Mexico; 27 of these were described from the 2016 field season and were reported previously (Marroquin-Flores et al. 2017). In that study, the total expected parasite richness for these communities was estimated to be ∼70 (95% CI: 43−98) haplotypes. When we consider only the same 49 host species that were sampled in 2016 and included in the previous haplotype diversity predictions, we detected 96 parasite haplotypes in the 2017 host samples. This empirical result falls at the upper end of this predicted richness. Of the three mountain ranges, Jemez had the highest parasite richness (61 haplotypes) despite having the fewest birds sampled. We detected 53 haplotypes in Mt. Taylor and 47 haplotypes in Zuni.

### Phylogenetic relationships and host specialization

The three haemosporidian genera were monophyletic with high posterior support, although relationships within each genus were less certain (Fig. 2). *Parahaemoproteus* and *Leucocytozoon* were detected in all host taxonomic groups sampled; *Plasmodium* was detected in all groups except non-passerines. Host breadth ranged from one to 14 hosts utilized, and MPD ranged from 0.9 to 20.7 with an average MPD of 10.2. Haemosporidians determined to be specialists were distributed across the parasite phylogeny, but they were most common in the genus *Parahaemoproteus*. Based on the SES_MPD_ metric, we classified 16 haemosporidians as specialists: 11 *Parahaemoproteus*, one *Plasmodium*, and four *Leucocytozoon* (Table 1). To assess community turnover between specialist and generalist parasites, we assigned the 16 most generalized parasites by considering those with the highest SES_MPD_ values. These included four *Parahaemoproteus*, five *Plasmodium*, and seven *Leucocytozoon*.

**Table 1.** Host specificity indices for haemosporidian haplotypes sampled at least twice (n ≥ 2). MPD is the weighted mean pairwise distance calculated in the ‘picante’ R package. The standardized effect size of MPD (SES_MPD_) and P-values were estimated using 999 randomizations. * indicates significantly specialized parasites (P < 0.05 or sampled 4 or more times from a single host).

| Haplotype | N | Host species | MPD | $SES_{MPD}$ | P-value | Classified as: |
| --- | --- | --- | --- | --- | --- | --- |
| H_PACPEC02 | 6 | 3 | 5.667 | -2.266 | 0.019* | Specialist |
| H_PIRLUD02 | 8 | 1 | NA | NA | * | Specialist |
| H_PIRFLA01 | 6 | 3 | 11.222 | -0.547 | 0.259 |  |
| H_SETAUD14 | 2 | 2 | 13.000 | 0.911 | 0.813 | Generalist |
| H_SPIPAS01 | 14 | 5 | 8.980 | -2.689 | 0.011* | Specialist |
| H_SPIPAS02 | 8 | 3 | 8.000 | -1.618 | 0.070 |  |
| H_CHOGRA01 | 5 | 1 | NA | NA | * | Specialist |
| H_JUHYE03 | 2 | 2 | 6.000 | -0.974 | 0.189 |  |
| H_PIRLUD01 | 6 | 1 | NA | NA | NA* | Specialist |
| H_TROAED12 | 15 | 3 | 6.738 | -1.859 | 0.060 |  |
| H_VIRPLU04 | 10 | 4 | 12.160 | -0.923 | 0.177 |  |
| H_VIRPLU01 | 17 | 5 | 10.228 | -2.185 | 0.033* | Specialist |
| H_VIGIL07 | 3 | 2 | 0.889 | -2.543 | 0.013* | Specialist |
| H_SIAMEX01 | 4 | 1 | NA | NA | * | Specialist |
| H_TABI02 | 2 | 2 | 2.000 | -2.145 | 0.048* | Specialist |
| H_GYMCYA02 | 3 | 2 | 0.889 | -2.348 | 0.012* | Specialist |
| H_SPISAL01 | 4 | 2 | 8.250 | -0.418 | 0.297 | Generalist |
| H_TACTHA02 | 5 | 1 | NA | NA | * | Specialist |
| H_EMPOBE02 | 2 | 2 | 4.000 | -1.621 | 0.099 |  |
| H_CYASTE05 | 4 | 2 | 9.750 | -0.060 | 0.393 | Generalist |
| H_PIRLUD07 | 3 | 2 | 13.333 | 0.974 | 0.879 | Generalist |
| P_MOLATE01 | 6 | 5 | 16.778 | 0.235 | 0.554 | Generalist |
| P_SEIAUR01 | 3 | 3 | 12.000 | -0.348 | 0.314 | Generalist |
| P_BAERID01 | 2 | 2 | 1.000 | -2.457 | 0.025* | Specialist |
| P_TACTHA01 | 3 | 3 | 11.111 | -0.662 | 0.237 |  |
| P_TACRUB04 | 2 | 2 | 14.000 | 1.215 | 0.939 | Generalist |
| P_PADOM09 | 3 | 3 | 17.778 | 1.348 | 0.927 | Generalist |
| P_TROAED24 | 18 | 8 | 17.148 | -0.531 | 0.274 | Generalist |
| P_LAIRI01 | 30 | 14 | 18.427 | -1.286 | 0.098 |  |
| L_CB1 | 7 | 4 | 14.367 | -0.226 | 0.359 | Generalist |
| L_CYASTE07 | 4 | 1 | NA | NA | * | Specialist |
| L_PIPMAC02 | 4 | 3 | 9.250 | -1.265 | 0.116 |  |
| L_SETGRA02 | 9 | 2 | 6.222 | -1.006 | 0.180 |  |
| L_PIRLUD03 | 5 | 2 | 0.960 | -2.354 | 0.012* | Specialist |
| L_VIGIL08 | 3 | 3 | 16.889 | 1.044 | 0.862 | Generalist |
| L_APHCAL01 | 6 | 4 | 14.222 | -0.281 | 0.344 | Generalist |
| L_COLBF21 | 22 | 11 | 19.289 | -0.006 | 0.488 | Generalist |
| L_SETNIG01 | 8 | 8 | 20.688 | 1.197 | 0.894 | Generalist |
| L_GYMCYA01 | 6 | 4 | 15.667 | 0.155 | 0.511 | Generalist |
| L_VIRPLU05 | 4 | 1 | NA | NA | * | Specialist |
| L_CNEORN01 | 3 | 2 | 7.111 | -0.863 | 0.181 |  |
| L_SETAUD25 | 2 | 2 | 8.000 | -0.515 | 0.250 | Generalist |
| L_PIRFLA02 | 5 | 3 | 6.880 | -1.940 | 0.048* | Specialist |

### Parasite community turnover

More than half of the parasite haplotypes were unique to a single mountain range. Of the 99 haplotypes, 56 were detected in one mountain range, 24 were shared among two mountain ranges, and only 19 were shared among all three mountain ranges. Comparing the observed data to the communities simulated under a model of no turnover indicated that there were more unique, single-mountain haplotypes (p < 0.001) and fewer haplotypes shared among all three mountains than expected by chance (p = 0.027; Fig. 3a). Turnover values between Mt. Taylor and the other two mountain ranges were higher than expected (Jemez-Mt. Taylor: p = 0.005, Mt. Taylor-Zuni: p = 0.01; Fig. 3b). The pruned dataset with identical host species indicated somewhat lower rates of turnover, with more unique haplotypes than expected (p = 0.013; Fig. 3c), no difference between the number of haplotypes shared among all three mountains (p = 0.084), and higher than expected turnover between Mt. Taylor and Jemez (p = 0.019; Fig. 3d).

**Figure 3.**
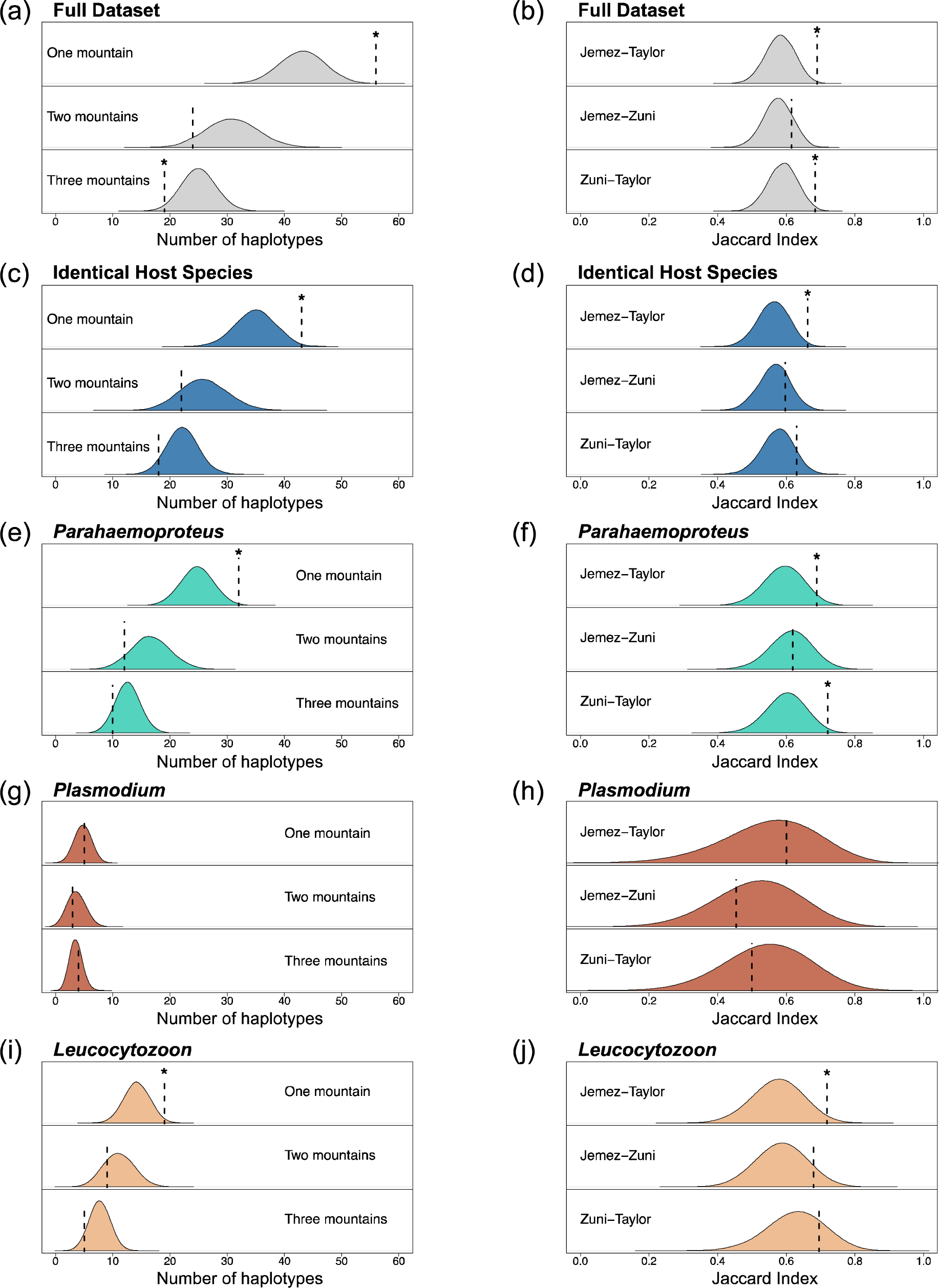
Null model results for full dataset (a,b), pruned dataset with identical host species (c,d), and the three parasite genera (e-j). Distributions are density plots from 10,000 simulations and summarize the number of haplotypes found in one, two, or three mountains (a,c,e,g,i) or the Jaccard indices between each pair of mountains (b,d,f,h,j). Dotted lines indicate observed values for each test. * indicates the observed value is significantly different (p < 0.05) from the simulated communities, assessed as the proportion of simulated values that are greater than, less than, or equal to the observed value.

Results for *Parahaemoproteus* and *Leucocytozoon* were similar to the overall dataset. We found more single-mountain haplotypes than expected (*Parahaemoproteus*: p = 0.007, *Leucocytozoon*: p = 0.023; Fig. 3e,i) and higher turnover indices than expected by chance between Mt. Taylor and Jemez (*Parahaemoproteus*: p = 0.037, *Leucocytozoon*: p = 0.019; Fig. 3f,j). For *Parahaemoproteus*, Mt. Taylor and Zuni were also more different than expected (p = 0.011). We found no significant differences between observed and expected communities of *Plasmodium* (Fig. 3g,h). The pruned genus-level datasets indicated an excess of one-mountain haplotypes for *Parahaemoproteus* (p = 0.032) and higher turnover than expected between Mt. Taylor and Jemez (p = 0.039; Online Resource 2; Fig. A1).

Parasites classified as specialists or generalists, when analyzed separately, showed no significant differences between the observed and simulated communities (Table 2). The parasite communities of both migrant and resident host species, analyzed separately, showed similar results as the overall community dataset. We found more single-mountain haplotypes than expected (migrant: p = 0.007, resident: p = 0.009), higher turnover than expected between Mt. Taylor and Zuni for migrant hosts (p = 0.04), and higher turnover than expected between Mt. Taylor and Jemez for resident hosts (p = 0.013; Table 2). Most of the focal host species showed no significant differences between observed and simulated communities, save for one of the best sampled hosts, *Vireo plumbeus*, which harbored fewer haplotypes that were shared between two mountain ranges than expected (p = 0.023; Online Resource 2; Fig. A2). The abundance distribution of range-restricted (single-mountain) haplotypes in the observed community indicated that at least some of these haplotypes were truly range-restricted because, rather than simply having low regional abundance, they were found 2–5 times on a single mountain, a phenomenon that was rarely observed in the simulated data (Online Resource 2; Fig. A3).

**Table 2.**
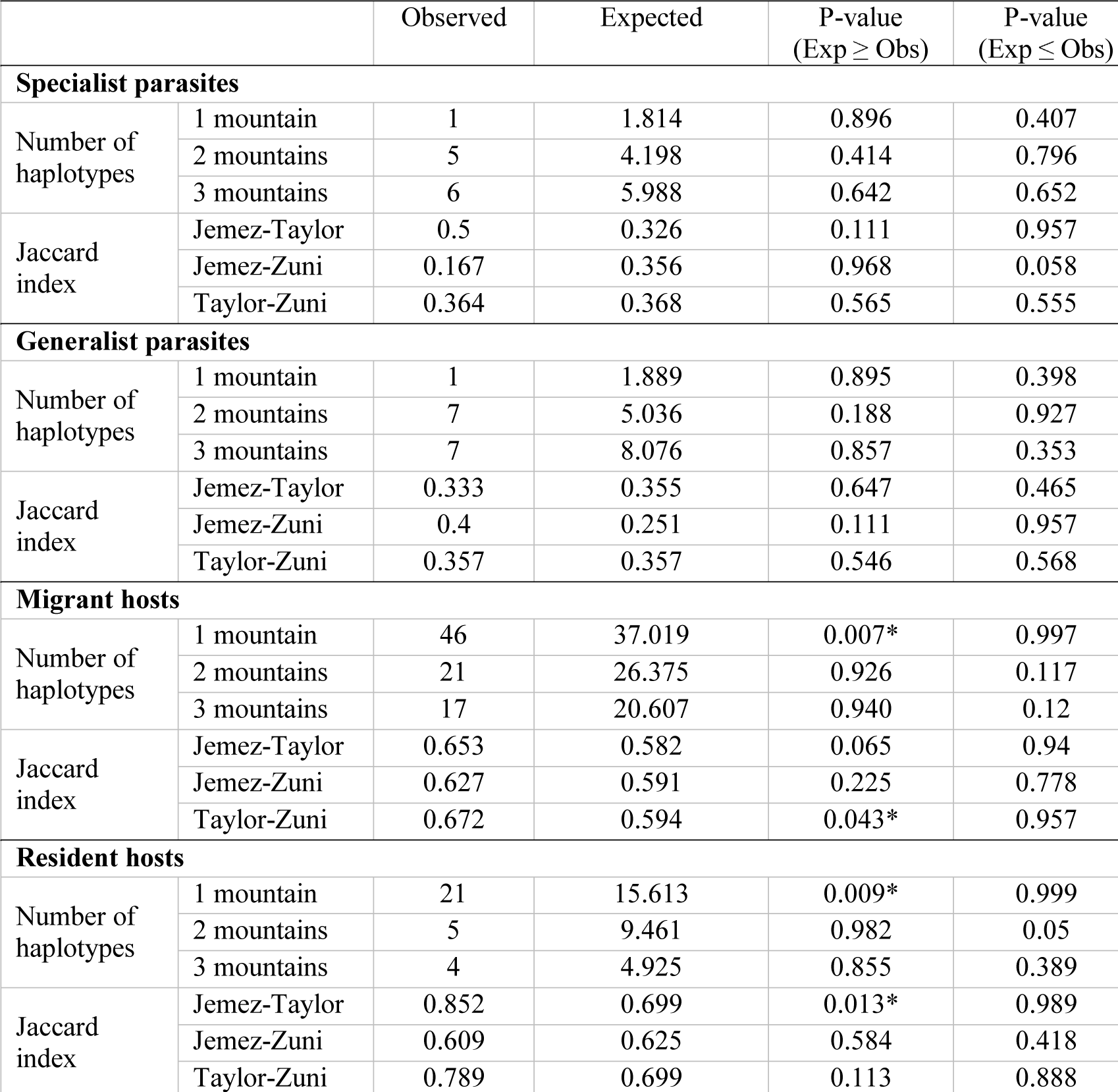
Null model results for specialist or generalist parasites, and migrant or resident host species. The number of haplotypes observed in a single mountain or shared between two or three mountains, and the Jaccard indices between each pair of mountains are compared to the expected values from 10,000 simulated communities. P-values were calculated as the proportion of simulated values that are greater than, less than, or equal to the observed.

| | | Observed | Expected | P-value<br>(Exp ≥ Obs) | P-value<br>(Exp ≤ Obs) |
| --- | --- | --- | --- | --- | --- |
| **Specialist parasites** |  |  |  |  |  |
| Number of haplotypes | 1 mountain | 1 | 1.814 | 0.896 | 0.407 |
|  | 2 mountains | 5 | 4.198 | 0.414 | 0.796 |
|  | 3 mountains | 6 | 5.988 | 0.642 | 0.652 |
| Jaccard index | Jemez-Taylor | 0.5 | 0.326 | 0.111 | 0.957 |
|  | Jemez-Zuni | 0.167 | 0.356 | 0.968 | 0.058 |
|  | Taylor-Zuni | 0.364 | 0.368 | 0.565 | 0.555 |
| **Generalist parasites** |  |  |  |  |  |
| Number of haplotypes | 1 mountain | 1 | 1.889 | 0.895 | 0.398 |
|  | 2 mountains | 7 | 5.036 | 0.188 | 0.927 |
|  | 3 mountains | 7 | 8.076 | 0.857 | 0.353 |
| Jaccard index | Jemez-Taylor | 0.333 | 0.355 | 0.647 | 0.465 |
|  | Jemez-Zuni | 0.4 | 0.251 | 0.111 | 0.957 |
|  | Taylor-Zuni | 0.357 | 0.357 | 0.546 | 0.568 |
| **Migrant hosts** |  |  |  |  |  |
| Number of haplotypes | 1 mountain | 46 | 37.019 | 0.007* | 0.997 |
|  | 2 mountains | 21 | 26.375 | 0.926 | 0.117 |
|  | 3 mountains | 17 | 20.607 | 0.940 | 0.12 |
| Jaccard index | Jemez-Taylor | 0.653 | 0.582 | 0.065 | 0.94 |
|  | Jemez-Zuni | 0.627 | 0.591 | 0.225 | 0.778 |
|  | Taylor-Zuni | 0.672 | 0.594 | 0.043* | 0.957 |
| **Resident hosts** |  |  |  |  |  |
| Number of haplotypes | 1 mountain | 21 | 15.613 | 0.009* | 0.999 |
|  | 2 mountains | 5 | 9.461 | 0.982 | 0.05 |
|  | 3 mountains | 4 | 4.925 | 0.855 | 0.389 |
| Jaccard index | Jemez-Taylor | 0.852 | 0.699 | 0.013* | 0.989 |
|  | Jemez-Zuni | 0.609 | 0.625 | 0.584 | 0.418 |
|  | Taylor-Zuni | 0.789 | 0.699 | 0.113 | 0.888 |

## Discussion

Here, we screened hundreds of birds for haemosporidian parasites across three mountain ranges in northern New Mexico. We demonstrated that avian haemosporidian communities in northern New Mexico sky islands are highly diverse and differ over relatively fine spatial scales, even when the same set of host species and habitats are sampled. By increasing the sampling from a previous survey of these three mountain ranges four-fold, we elaborated general patterns of parasite abundance, diversity, specialization, and variation in infection rate among host species in this system. Our null model tests revealed an excess of localized (single mountain) parasite haplotypes and a deficit of shared haplotypes relative to expectations under a homogeneous regional community. These null models demonstrated that apparent turnover between mountain ranges is real in some cases, and not merely an artifact of undersampling. In other cases, undersampling can clearly create the appearance of turnover when there is none, and it can also exaggerate the degree of turnover as indicated by common metrics.

Improved sampling in this system, from 186 birds in Marroquin-Flores et al. (2017) to 776 birds screened in the present study, showed that while parasite abundance remained highly consistent, parasite diversity had been underestimated. Overall prevalence (proportion of infected birds out of total screened) was 36.6% in 2016 and 36.1% in the combined-year dataset. Prevalence estimates for parasite genera were also consistent: *Parahaemoproteus* was estimated at 20.9%, and now at 20%; *Plasmodium* remained 8%, and *Leucocytozoon* was estimated at 13.4%, and now at 13.5%. The number of co-infections detected increased slightly in the combined-year dataset, from 9.7% to 11.6% of birds screened. *Parahaemoproteus* was the most common genus in all three mountain ranges, and consistently had the highest richness (54 haplotypes, 55% of total), while *Plasmodium* had the lowest (12 haplotypes, 12% of total). Total parasite richness increased from 43 haplotypes detected in 2016 to 99 haplotypes total, which was higher than extrapolated diversity estimates (∼70 haplotypes, 95% CI: 43–98) from the previous survey of these three mountain ranges (Marroquin-Flores et al. 2017). When only the 49 host species sampled in 2016 were considered, we found 96 haplotypes, indicating that the addition of new host species alone does not account for this higher than expected diversity. Instead, a likely explanation is that true turnover between sites (beta diversity) exists and is an unaccounted for source of variation affecting these estimates, each of which was derived from the combined communities of the three mountains. When we extrapolate diversity from the full dataset, we continue to predict that a high proportion (∼50%) of diversity in this system has yet to be discovered (∼151 haplotypes, 95% CI: 115–188). These results indicate that extrapolated measures of diversity are likely to be underestimates when structured communities are analyzed together, or when there are any unmodeled sources of variation such as heterogeneous habitats or host communities.

Our results refined several interesting patterns related to variation in host breadth among parasites. Previously, *Parahaemoproteus* and *Leucocytozoon* were detected from only three of six host clades (Marroquin-Flores et al. 2017); here we found infections in all host clades, although both genera were still most common in Corvides and Passerides 1b (Fig. 2). Each of the three genera of haemosporidian parasites contained haplotypes that we classified as specialists and generalists, respectively, although *Parahaemoproteus* included the most host specialist lineages and *Plasmodium* included the fewest. In previous work, clades of haemosporidian haplotypes associated with vireos have been noted in the western U.S. (Walther et al. 2016; Marroquin-Flores et al. 2017) and the eastern U.S. (Ricklefs et al. 2005). One outstanding question relates to whether these putative ‘vireo specialist’ parasites can establish successfully in other host species. Here we found that three of the 12 ‘vireo’ haplotypes did occur in other host clades, at least occasionally (Fig. 2). Only one of these infections out of eight screened by microscopic examination was confirmed to contain gametocytes. In this case, the parasite was VIRPLU04 and the host was a warbler, *Setophaga nigrescens*. This indicates that at least one of these vireo associated parasites is able to establish and reproduce in a non-vireo host.

Increased sampling also clarified patterns of variation in prevalence among host species. Previously, no infections in non-passerines or certain passerine species, such as nuthatches, were found in this system. While non-passerines were still rarely infected, we detected infections in six of 51 birds screened (11.8%) and three of these were confirmed by microscopic examination. Nuthatches also had very low prevalence; we did detect infections in five of 80 birds (6.25%) by PCR, but found no gametocytes in thin blood smears, suggesting that these might represent abortive infections of atypical hosts (Olias et al. 2011; Moens et al. 2016). At the clade level, suboscines were the least infected, a pattern that has been demonstrated consistently and has been used to illustrate that susceptibility to haemosporidians is evolutionarily conserved (Ricklefs 1992; Barrow et al. 2019). Corvides was the most infected host clade, with 58 of 78 birds (74%) infected and a total of 97 infections detected. Vireos were among the host species with the highest prevalence; 87% of *Vireo plumbeus* and 78% of *V. gilvus* individuals were infected. Although sampling was limited (n=6), 100% of Woodhouse’s Scrub-Jay (*Aphelocoma woodhouseii*) individuals were infected and five of six were co-infected with *Leucocytozoon* and one of the other two genera. Similarly high prevalences have been observed in *V. gilvus* in California (Walther et al. 2016) and in European corvids including crows and magpies (Scaglione et al. 2016; Schmid et al. 2017).

Null models were useful for detecting community differences while overcoming sampling inadequacies that are inevitable in complex, host-parasite systems. We found more unique parasite haplotypes (occurring in a single mountain) and fewer haplotypes shared across all three mountains than expected under a model of no turnover. When we pruned the dataset to include only identical host species across mountain ranges, these patterns were somewhat weaker, statistically, but consistent. Host community turnover is expected to play a role in parasite community turnover (Clark et al. 2018; Williamson et al. 2019). By effectively controlling for the host community, we demonstrate that haemosporidian parasite communities even differ among sky islands when the same set of host species are included. When considering the three genera separately, we found that both *Parahaemoproteus* and *Leucocytozoon* had similar patterns with the overall community, with more unique, single-mountain haplotypes than expected. *Plasmodium* did not show significant turnover patterns and, although this genus is rare in this system, this negative result was consistent with the expectation that *Plasmodium* haplotypes tend to be generalized and broadly distributed (e.g., Hellgren et al. 2015). Parasite turnover on fine spatial scales, such as between adjacent sky islands, is important because it implies that a single host species will encounter distinct parasite assemblages and associated diversifying selective pressures across its range.

Our results were consistent with previous studies demonstrating that geographic distance was not a consistent predictor of haemosporidian parasite turnover (Ellis et al. 2015; Williamson et al. 2019). Interestingly, the centrally-located mountain range in this system, Mt. Taylor, exhibited the highest degree of turnover with the other two mountain ranges. These sampled communities are separated by a straight-line geographic distance of only ∼50–100 km. Community differences on this fine scale are not unprecedented; Williamson et al. (2019) found high turnover in the haemosporidians of Audubon’s Warblers (*Setophaga auduboni*) sampled from eight mountain ranges in the southwestern U.S., including the three in the present study. The haemosporidian communities sampled in the Jemez and Zuni Mountains, separated by a distance of ∼160 km, did not differ more than expected, a result that would have been difficult to interpret from a traditional beta diversity metric (Jaccard dissimilarity index: 0.615). Null model tests thus provided additional insights into the patterns of turnover in this system. The lack of the distance-decay relationship typical of some parasite systems (Poulin 2003; Thieltges et al. 2009) has several possible explanations. Subtle habitat differences leading to changes in the vector community, which remains to be sampled and characterized, could influence some of the differences in parasite communities. The entire avian community, including host species not sampled in this study, also differs slightly between mountain ranges and partly explains parasite community turnover (Williamson et al. 2019). We suggest it is most likely that the turnover patterns we observed represent stochastic colonization-extirpation dynamics on sky islands.

We further explored these results by testing whether haemosporidian communities from migrant or resident hosts differed in patterns of turnover. Both migrant and resident host species had similar patterns compared to the overall host community, with more unique haplotypes than expected by chance, and community differences between Mt. Taylor and the other two mountain ranges. Long-distance migrants are expected to connect avian haemosporidian communities in breeding and wintering grounds, and greater connectivity between parasite faunas has been documented in the Americas (eastern North America and West Indies/northern South America) compared to the Euro-African migration system (Hellgren et al. 2007; Ricklefs et al. 2017). Our results suggest that on a regional scale, migration does not strongly affect spatial community structure of avian haemosporidians, at least at the spatial scale of adjacent sky islands.

Here, we established a framework to compare communities in which sampling is inevitably incomplete and subject to stochastic variation. While accounting for several common sampling issues — skewed parasite-haplotype abundance distributions, uneven sampling among sites, different sets of host species, and habitats — we confirmed that avian haemosporidian turnover occurs on fine spatial scales, dispersal barriers of tens of kilometers, across which host turnover is negligible. Importantly, we were able to build on the work of Marroquin-Flores et al. (2017) because all data are available via museum specimens and open-access databases (Arctos, GenBank, and MalAvi). Likewise, future studies can continue building on these datasets to compare avian haemosporidian assemblages across mountain ranges, elevational zones, host communities, and focal host species. The differences we have already detected among nearby sites with sampling from the same habitats, elevations, and host communities suggest that adding sites with further habitat, climate, and host variation will provide rich datasets for understanding the drivers of diversity in this complex system.

## Acknowledgements

We thank Michael Andersen, Sara Brant, Mariel Campbell, Joseph Manthey, Moses Michelsohn, and George Rosenberg. This work was supported by the Bureau of Land Management Rio Puerco Field Office (via the Colorado Plateau Cooperative Ecosystems Studies Unit agreement) and NSF DEB-1146491. SMB, RMF, and TEM were supported by PREP/FlyBase Fellowships (NIH 5R25HG007630) and LNB was supported by an NSF Postdoctoral Research Fellowship in Biology (NSF DBI-1611710).

