## Supplementary Figures for "Detecting turnover among complex communities using null models: A case study with sky-island haemosporidian parasites"

**Electronic Supplementary Material**

**Appendix 2**

Figure A1. Null model results for pruned datasets for the three parasite genera.

Figure A2. Null model results for focal host species.

Figure A3. Comparison of range-restricted haplotype abundance distributions.

**Figure A1.** Null model results for pruned datasets for the three parasite genera. Distributions are density plots from 10,000 simulations and summarize the number of haplotypes found in one, two, or three mountains (a,c,e) or the Jaccard indices between each pair of mountains (b,d,f). Dotted lines indicate observed values for each test. * indicates the observed value is significantly different (p < 0.05) from the simulated communities, assessed as the proportion of simulated values that are greater than, less than, or equal to the observed value.

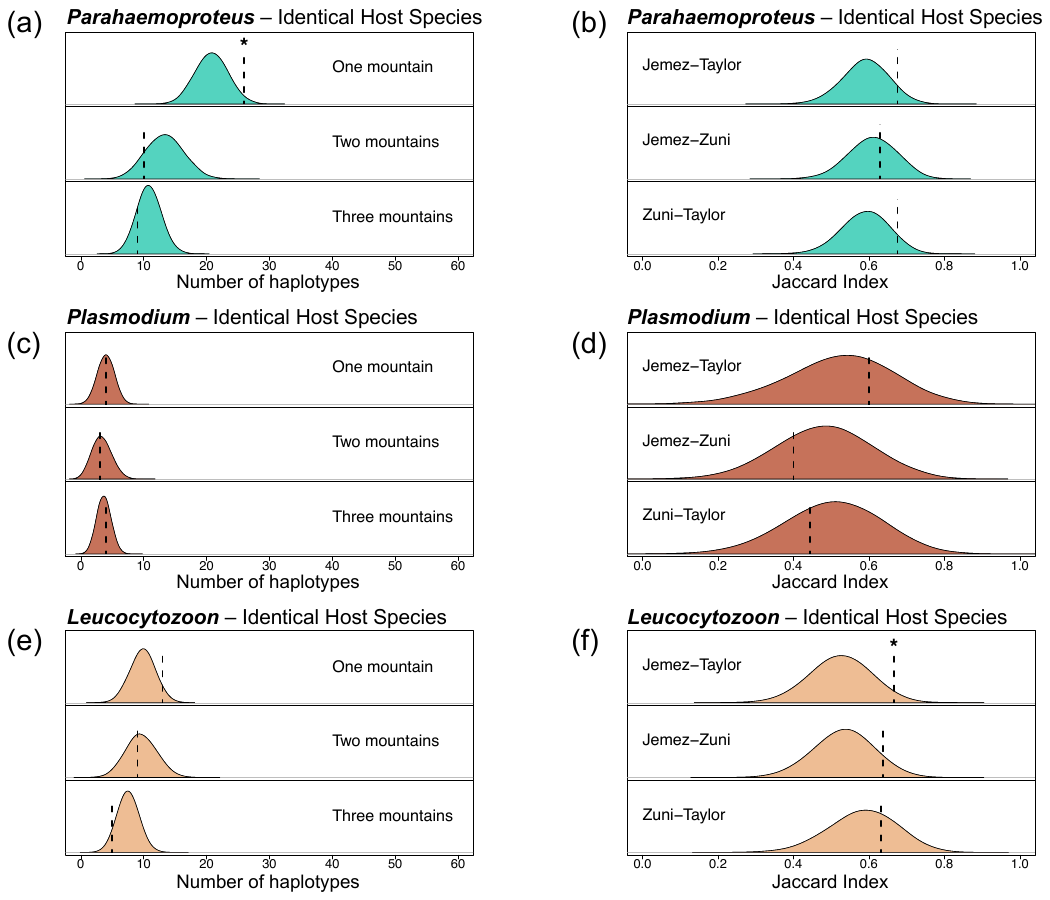

**Figure A2.** Null model results for focal host species. Distributions are density plots from 10,000 simulations and summarize the number of haplotypes found in one, two, or three mountains (a,c,e) or the Jaccard indices between each pair of mountains (b,d,f). Dotted lines indicate observed values for each test. * indicates the observed value is significantly different (p < 0.05) from the simulated communities, assessed as the proportion of simulated values that are greater than, less than, or equal to the observed value.

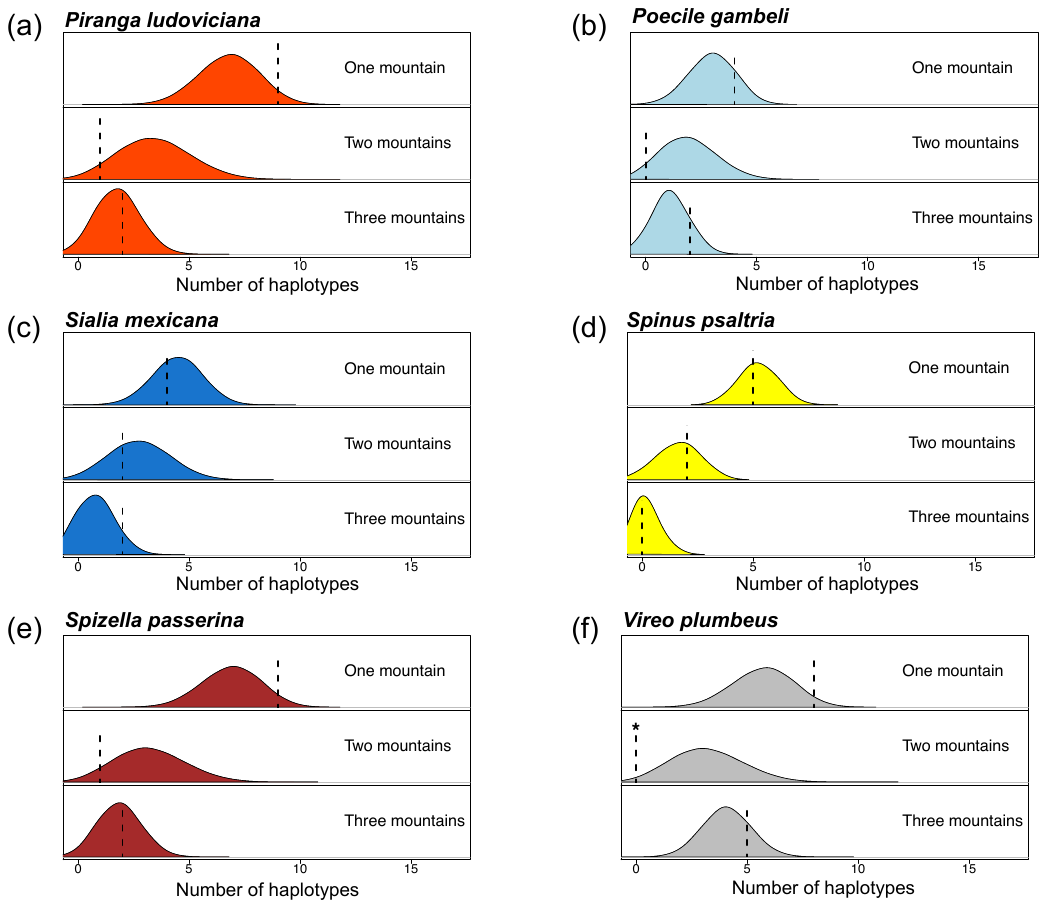

**Figure A3.** Comparison of range-restricted (single mountain) haplotype abundance distributions for observed (red line) and 10,000 simulated (light grey) communities. The black line is the average of the simulated communities. We predicted that range-restricted haplotypes in the simulated communities would be dominated by rare haplotypes (haplotype abundance = 1) because haplotypes are assumed to be distributed throughout the region. The abundance distribution for the observed community should be slightly right-shifted if at least some range-restricted haplotypes have higher abundances, indicating they are truly range restricted.

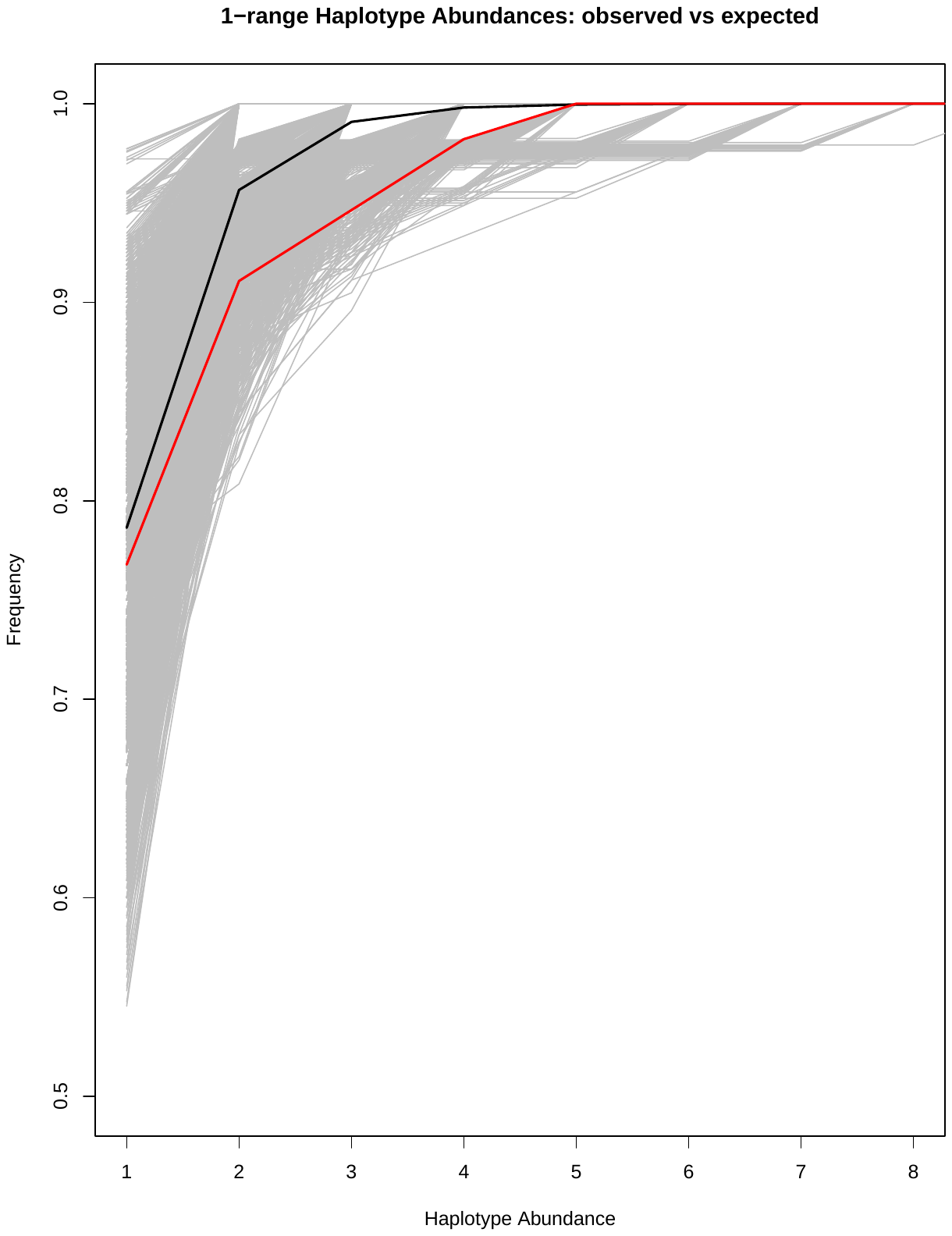
